# Humanized *CYP2C19* transgenic mouse as an animal model of cerebellar ataxia

**DOI:** 10.1101/2022.01.10.475612

**Authors:** Filip Milosavljević, Irene Brusini, Andrea Atanasov, Marina Manojlović, Maria Novalen, Marija Vučić, Zorana Oreščanin Dušić, Jelena Brkljačić, Sharon Miksys, Čedo Miljević, Aleksandra Nikolić, Duško Blagojević, Chunliang Wang, Peter Damberg, Vesna Pešić, Rachel F Tyndale, Magnus Ingelman-Sundberg, Marin M Jukić

**Affiliations:** Department of Pharmacology, Faculty of Pharmacy, University of Belgrade, Serbia; Department of Biomedical Engineering and Health Systems,KTH Royal Institute of Technology, Huddinge, Sweden; Department of Neurobiology, Care Sciences and Society, Karolinska Institute, Huddinge, Sweden; Campbell Family Mental Health Research Institute, Centre for Addiction and Mental Health, Toronto, Ontario, Canada; Department of Psychiatry, University of Toronto, Ontario, Canada; Department of Pharmacology and Toxicology, University of Toronto, Ontario, Canada; Institute for Biological Research “Siniša Stanković”, Belgrade, Serbia; Department of Psychiatry, Faculty of Medicine, University of Belgrade, Serbia; Institute for Mental Health, Belgrade, Serbia; Karolinska Experimental Research and Imaging Center, Karolinska University Hospital, Solna, Sweden; Section of Pharmacogenetics, Department of Physiology and Pharmacology, Karolinska Institute, Stockholm, Sweden

## Abstract

**Background:** Animal models are essential for understanding etiology and pathophysiology of movement disorders. Previously, we have found that mice transgenic for the human *CYP2C19* gene, expressed in the liver and developing brain, exhibit altered neurodevelopment associated with impairments of their motor function and emotionality.

**Objectives:** To characterize motoric phenotype of the *CYP2C19* transgenic mice and validate its usefulness as an animal model of ataxia.

**Methods:** The rotarod and beam-walking tests were utilized to quantify the functional alterations induced by motoric phenotype. Dopaminergic system was assessed by tyrosine hydroxylase immunohistochemistry and by chromatographic quantification of the whole-brain dopamine levels. Beam-walking test was also repeated after the treatment with the dopamine receptor antagonists, ecopipam and raclopride. The volumes of 20 brain regions in the CYP2C19 transgenic mice and controls were quantified by 9.4T gadolinium-enhanced *postmortem* structural neuroimaging.

**Results:** *CYP2C19* transgenic mice were found to exhibit abnormal, unilateral ataxia-like gait, clasping reflex and 5.6-fold more paw-slips using the beam-walking test (*p*<0.0001, n=89); the phenotype was more pronounced in younger animals. Hyperdopaminergism was observed in the *CYP2C19* mice; however, the motoric impairment was not ameliorated by dopamine receptor antagonists and there was also no midbrain dopamine neuron loss in *CYP2C19* mice. However, in these mice, cerebellar volume was drastically decreased (-11.8% [95%CI: −14.7, −9.0], *q*<0.0001, n=59), whereas a moderate decrease in hippocampal volume was observed (−4.2% [95%CI: −6.4%, −1.9%], *q*=0.015, n=59).

**Conclusions:** Humanized *CYP2C19* transgenic mice exhibit altered motoric function and functional motoric impairments; this phenotype is likely caused by an aberrant cerebellar development.

## INTRODUCTION

Cerebellar ataxia is an umbrella clinical term used for the group of severe illnesses caused by various factors, ranging from hereditary genetic mutations to infections, intoxications and degenerative diseases^1^. Clinical phenotype can be heterogeneous between different types of cerebellar ataxia; the most common symptoms include unstable gait, lack of balance, blurred vision, slurred speech and a loss of fine dexterity^1^. Cerebellar ataxia has a major negative impact on the quality of life in affected individuals; however the etiology and pathophysiology of the disease is still not completely understood and effective treatment strategies for many forms of ataxia are sparse^2,3^.

Several animal models of cerebellar ataxia have been used in the past in an attempt to clarify molecular and cellular mechanisms behind cerebellar function impairments. For certain models of ataxia, specifically engineered transgenic mice that carry gene variants found in human hereditary forms of ataxia are used; while other rodent models are selected for their ataxia-like phenotype by selective breeding^4^. Since the animal models have been very useful in the research of cerebellar functions and ataxia pathophysiology in the past^2^, characterization of novel animal models of ataxia clearly holds a potential to further improve the understanding of the cerebellar phenotypes.

CYP2C19 is an enzyme expressed mainly in the liver; however the expression is also detected in fetal life in the brain, meaning that the enzyme could be involved in neuronal development^5,6^. *CYP2C19* transgenic mouse carries the human *CYP2C19* gene, which is not present in a mouse genome^7^. The mice do not have an orthologous variant of *CYP2C19* and the transgenic mice exhibit the same CYP2C19 enzyme organ distribution and expression during development as seen in humans^5^. Therefore, this mouse model is a valid tool for studies of the role of the CYP2C19 enzyme in developing brain and its potential involvement in neurodevelopment. Previous research on *CYP2C19* mice described their complex emotional phenotype that includes increased susceptibility to stress^6^, elevated depression like behavior^5^, abnormal serotoninergic neurotransmission^5^, and decreased whole-brain and hippocampal volumes^6^; in addition, alterations in motoric functions have also been observed^6^.

The initial aim of this study was to in-depth characterize motoric phenotype in *CYP2C19* humanized transgenic mice, while the subsequent aim was to examine whether *CYP2C19* transgenic mouse can be validated as an animal model of cerebellar ataxia.

Primary hypothesis was: **(1)** *CYP2C19* transgenic mice exhibit ataxia-like motoric phenotype that causes functional motoric impairment; while the secondary hypotheses were: **(2)** Structural and functional alterations of *nigro-straiatal* dopaminergic system are involved in the observed motoric phenotype; **(3)** There are volumetric abnormalities in the brain regions of *CYP2C19* mice that are likely related to its complex motoric and emotional phenotype; **(4)** Oxidative stress and/or neuromelanin accumulation are involved in the structural changes in the brain.

## METHODS

### Laboratory Animals

The *CYP2C19* transgenic (TG) mice were generated from the C57Bl/6JOlaHsd genetic background and are hemizygous carriers of 12 copies of human *CYP2C19* and *CYP2C18* genes^7^. Animals were housed in groups of 3-5 animals per cage. Housing conditions included 12h light/dark cycle, controlled room temperature (22±1°C), humidity (40-70%) and illumination; and *ad libitum* access to water and pelleted food. Adult mice (>10 weeks) of both sexes were included, and they were divided into two test groups based on their genotype: *CYP2C19* transgenic mice and their wild type littermates (WT). Researchers conducting the experiments were blinded for animals' genotype whenever this was possible. General signs of animal wellbeing such as level of activity, fur condition, weight loss etc. were checked at least twice a week. If signs of severe distress were observed, such animals were excluded from the experiment and euthanized to reduce their suffering. Also, the animals showing abnormal behavior such as repetitive movements, high agitation, and lack of motivation to complete experimental tasks were excluded from the subsequent analysis. All experiments were done according to the permit of Ethical Committee on Animal Experimentation of the University of Belgrade – Faculty of Pharmacy, Serbia (permit number 23-07-00933/2019-05 to MJ). Study was conducted and reported in accordance with ARRIVE 2.0 guidelines^8^.

### Motor Tests

In CYP2C19 transgenic mice (TG mice), gait was visually analyzed in freely moving animals and compared to WT controls. Mice were also visually screened for pathological clasping reflex, which is present in many motorically impaired rodent strains^9^. Motoric function in mice was quantified by the rotarod and beam walking tests. In the rotarod test, Ugo Basile Rotarod 47700 (Ugo Basile, Italy) apparatus with four 8.7cm wide lanes was used with the accelerating rotarod protocol. Latency to falling was measured, and an average of the three longest runs was used as readout of motoric performance; since the motorically impaired mice tend to fall sooner than the animals with the normal motor function.

In the beam-walking (BW) test, rectangular 8 mm wide and 1 m long transparent acrylic beam was used. Since trained and motivated animals tend to cross the beam as fast as they possibly can, beam-crossing time is prolonged mainly due to the motoric impairment. Also, animals with motoric impairment tend to exhibit poorer motor coordination and more paw slips during the beam-crossing. Hence, the performance in the BW test was measured as the average of the three shortest beam-crossing times and as the average of three smallest measured numbers of paw-slips per run.

### Dopamine Concentration Determination and Dopamine Receptor Antagonist Treatment

Concentration of dopamine was determined in the brain hemispheres of 3 months old mice using HPLC-MS-MS method described elsewhere^10^. Noteworthy, this was the only experiment which included only males, since the females had to be spared for the breading and colony expanding purpose at the time this experiment was conducted.

To test the effects of dopaminergic receptor agonists on the motoric performance of TG mice, the BW test was repeated after the treatment with the selective D1 antagonist ecopipam (SCH-39166, Tocris Bioscience, UK) or the D2 antagonist raclopride (Tocris Bioscience, UK). Ecopipam hydrobromide was given in the dose of 0.1 mg/kg as a free base and raclopride was given at the dose of 0.25 mg/kg. Drugs were dissolved in the saline solution and administered to the animals intraperitoneally, 30 minutes before the BW test. Doses were selected to be sufficient to impact the performance of animals with hyperactivated dopaminergic system, but not to cause general sedation of the animals^11^.

### Immunohistochemistry and Neuromelanin Staining

Immunohistochemically stained slides of mouse brain sections were used to investigate the structures of mid-brain dopaminergic nuclei in mice. Tyrosine Hydroxylase positive (TH+) cells were counted in *substantia nigra pars compacta* (SN) and *ventral tegmental area* (VTA) (Staining validation presented in Figure 2C) and compared in 9 TG/WT littermate pairs; 4 pairs were adults (6-month-old), and 5 pairs were at the old age (15-month old). Nine representative coronal slides were analyzed and neuron was considered TH+ if the cytoplasm was stained and if the nucleus was visible.

The presence of neuromelanin was assessed in the brain sections of 15-month-old mice stained with Fontana-Masson melanin stain (ab150669, Abcam, UK) according to the manufacturer supplied protocol^12^. Neuromelanin is a side-product of catecholamine synthesis and is potentially involved in the pathogenesis of dopamine cell death in humans^13^. Hence, it was hypothesized that increased dopamine production in the brains of TG mice could be the cause of substantial aggregation of neuromelanin, and could be a sign of increased vulnerability of dopaminergic neurons to the cell death.

### Gd-enhanced Neuroimaging

Structural differences in brains of WT and TG mice were investigated with gadolinium (Gd) enhanced structural 9.4T MRI *ex-vivo* neuroimaging. The samples were prepared for ex-vivo MRI imaging through transcardial perfusion with formaldehyde, supplemented with the MRI contrast agent ProHance. Extra-cranial tissue was removed and four heads were wrapped in Gaze and placed in a 30 ml syringe. The void of the syringe was filled with perfluorinated oil, i.e. fomblin (Solaway Solexis), as a non-signaling susceptibility matched environment for the heads. The samples were scanned using a 9.4 T horizontal bore MRI scanner (Varian, Yarnton, UK) equipped with a millipede coil with an inner diameter of 30 mm. The samples were scanned using a multi gradient echo 3d with the following parameters: matrix size 1024×512×512, Field-of-View 51.2×25.6×25.6 mm^3^, recovery time 30 ms, time to echo 2.82 ms, flip angle 35°. Two echoes separated by 4.96 ms were acquired and 512 dummy excitations to establish steady state were used. Obtained brain images were segmented automatically based on the 3D Waxholm Space 2012 mouse brain atlas^14,15^ into 39 brain regions, but 19 regions such as ventricles, inner ear, medulla oblongata, various nerves and white matter structures were excluded from the analysis due to proneness of substantial variability induced by tissue processing. Volumes of each brain region were quantified by multiplying the number of voxels the region comprises with the volume of individual voxel (50 μm voxel edge, 12.5 x 10^-5^ mm^3^), while the total brain volume was calculated as the sum of volumes of all 39 brain regions.

Subsequently, between-group differences in local grey matter (GM) volume were further investigated by performing voxel-based morphometry. We employed FMRIB Software Library voxel-based morphometry (FSL-VBM)^16^, an optimized VBM protocol^17^ carried out with FSL tools^18^. Though, in our work, the protocol was slightly modified in order to adapt it to the present mouse brain images and to compensate for the big difference between human and mouse brains. Local differences between groups were investigated by calculating voxel-wise F statistics for genotype, using genotype and sex as factors.

### Anti-oxidative enzyme concentration and activity

To assess potential differences in the brain oxidative-antioxidative balance between the WT and TG mice, antioxidative enzyme status was determined in the brain tissue. Concentrations and activities of four antioxidative enzymes: superoxide dismutase (SOD), catalase (CAT), glutathione peroxidase (GPx) and glutathione reductase (GR) were measured in the mouse cerebellum, hippocampus and whole hemispheres. All enzyme activities were determined biochemically and expressed as units (U) per mg of protein, while enzyme quantities were determined with Western blot method and presented in arbitrary units.

### Statistics

Data was analyzed with SPSS statistics 20 software (IBM, USA). Genotype was the principle independent variable in all experiments. Criteria for the selections of sample sizes are presented in the supplementary material. Normality of data distributions was assessed with Shapiro-Wilk test; if distribution was normal all outliers were excluded based on the 2.2 IQR rule^19^. If experiment included covariates such as age or sex, 1-way ANCOVA was performed; otherwise, the Student's *t*-test for independent samples was used. Exceptions were experiments that included drug treatments where 2-way mixed ANCOVA was performed (treatment and genotype were used as independent variables), and non-normally distributed data where Mann Witney and Kuskall Wallis tests were used. All tests were two-tailed with *p-* value <0.05 as a measure of significance. False discovery rate (FDR) correction of *p*-values for multiple comparisons was applied when multiple brain regions and multiple antioxidant enzyme activities were analyzed. Differences between test groups were textually presented as the ratio-of-means with 95% confidence interval (95%CI) for normally distributed data, or as median and interquartile range (IQR) for each group for non-Gaussian data. All experimental protocols were pre-specified during the ethics committee permit acquirement process. For most of the experiments, no randomization technique was employed in the experimental design due to the nature of independent variable i.e. mutants were always compared with controls. For the beam-walking experiment which included drug treatment, the mutants and controls were randomized into treatment groups to ensure that the main readout (beam-crossing time) is equivalent in all experimental groups.

Detailed descriptions of all experimental procedures and statistical analyses are available in the supplementary materials.

## DATA SHARING

The data that supports the findings of this study are available in the supplementary material of this article

## RESULTS

Initially, the motoric phenotype of the transgenic *CYP2C19* (TG) mice was investigated visually; the TG mice exhibited altered gait, characterized by excessive hind paw elevation (Figure 1A, Video1). Also, pathological clasping reflex was observed, which occurred after several seconds of struggle (Figure 1A). Observed phenotype was more pronounced in young animals with higher paw elevation than in adults and short paw-locking in the uppermost position. Also, younger TG mice exhibited bilateral phenotype that became unilateral between 6^th^ and 10^th^ week of age (Video1). To investigate potential functional implications of the observed TG motoric phenotype, the latency to falling was measured on the rotarod test, while the beam-crossing time and number of paw slips were measured on beam-walking test. On the rotarod test, no significant motoric impairment was observed (*p*=0.93), as TG and wild type (WT) mice did not exhibit different latencies to falling (Figure 1B). However, impairment of motor function was observed on the beam walking test, as the TG mice exhibited slightly longer (1.14-fold, [95%CI: 1.06, 1.22], *p*=0.0014) beam crossing time and profoundly 5.6-fold increased (TG: 1.7 [IQR: 1.0, 2.3] vs WT 0.33 [IQR: 0.0, 0.92], p<0.0001) number of paw slips compared to WTs (Figure 1C). In summary, TG mice exhibit a visually pronounced motoric phenotype, which impacts their motoric function in challenging tasks.

**Figure 1:**
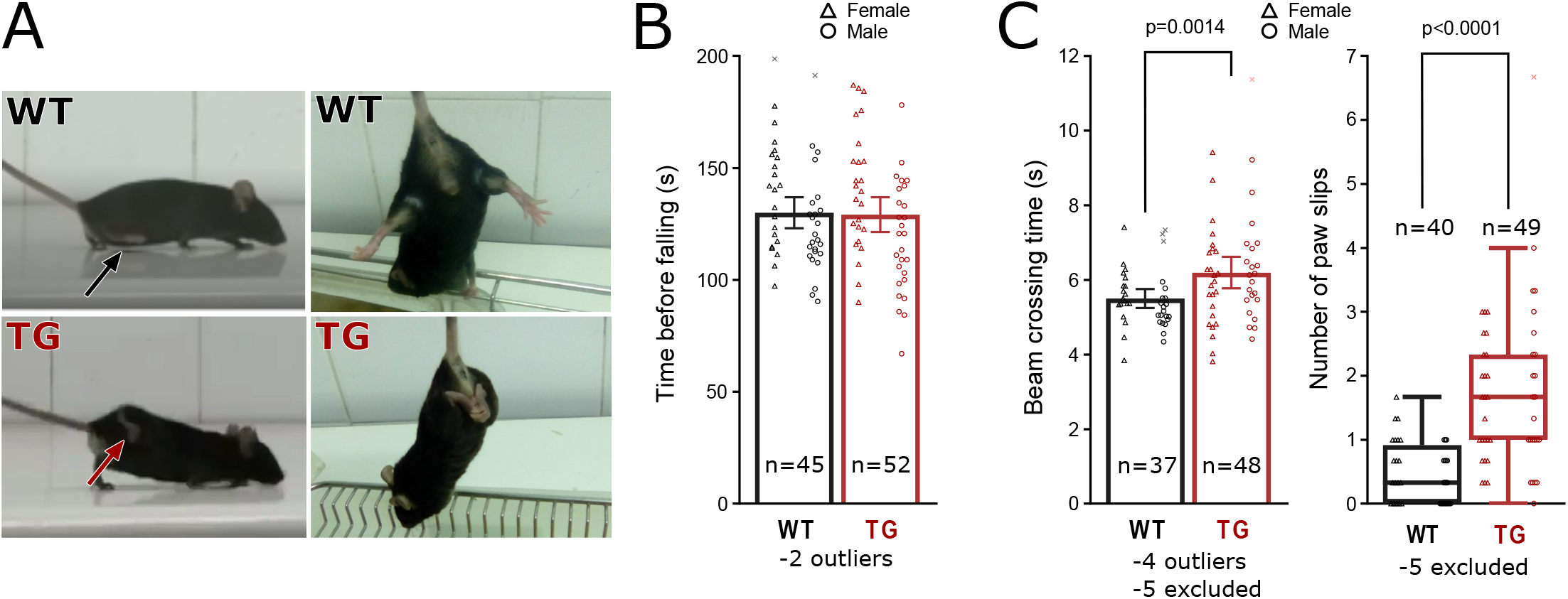
Motoric phenotype in TG mice. *CYP2C19* transgenic mice exhibit motor phenotype with altered gait **(A, left)**, clasping reflex **(A, right),** no change in the rotarod test **(B)** and worse performance in the beam-walking test **(C)**. *WT – Wild Type mice; TG – CYP2C19 transgenic mice.*

Next, the structure and function of dopaminergic system in TG mice was investigated; TG mice exhibited a 1.15-fold increase (95%CI: 1.09, 1.22; *p*<0.0001) in the whole brain dopamine concentration compared to WTs (Figure 2A). To investigate the effect of D1 antagonist ecopipam and D2 antagonist raclopride on the previously observed motor phenotype of TG mice, animals repeated beam walking test shortly after the acute drug treatments. Both drugs failed to ameliorate the motor function impairment of TG mice; moreover, they slightly prolonged beam crossing time in all mice compared to saline treated mice (Ecopipam: 1.2-fold [95%CI: 1.17, 1.30], p<0.0001; Raclopride: 1.2-fold [95%CI: 1.12, 1.25], p<0.0001), without genotype specific changes (Figure 2B). To investigate the potential changes in shape and microstructure of dopaminergic nuclei, number of tyrosine hydroxylase positive (TH+) neurons was quantified throughout rostro-caudal axis. Since the mouse age did not significantly affect the number of dopaminergic neurons and since it did not interact with genotype in the initial analysis, this covariate was excluded from the analysis, and 6- and 15-month-old mice were considered as one homogenous group for both genotypes. In the section −3.52 mm from bregma, the number of dopaminergic neurons in *Substantia nigra* was slightly, but significantly (1.15-fold [95%CI: 1.03, 1.27], *p*=0.037) reduced in the TG compared to WT mice (Figure 2D); however, this change did not remain significant after multiple comparisons corrections (*q*=0.51). In addition, neuromelanin staining was negative in dopaminergic neurons in the *substantia nigra* in all mice, unrelated to the age and genotype. In summary, TG mice exhibit hyperdopaminergism, which is unlikely directly associated with the observed motor phenotype and which is not associated with very pronounced structural changes within dopaminergic nuclei.

**Figure 2:**
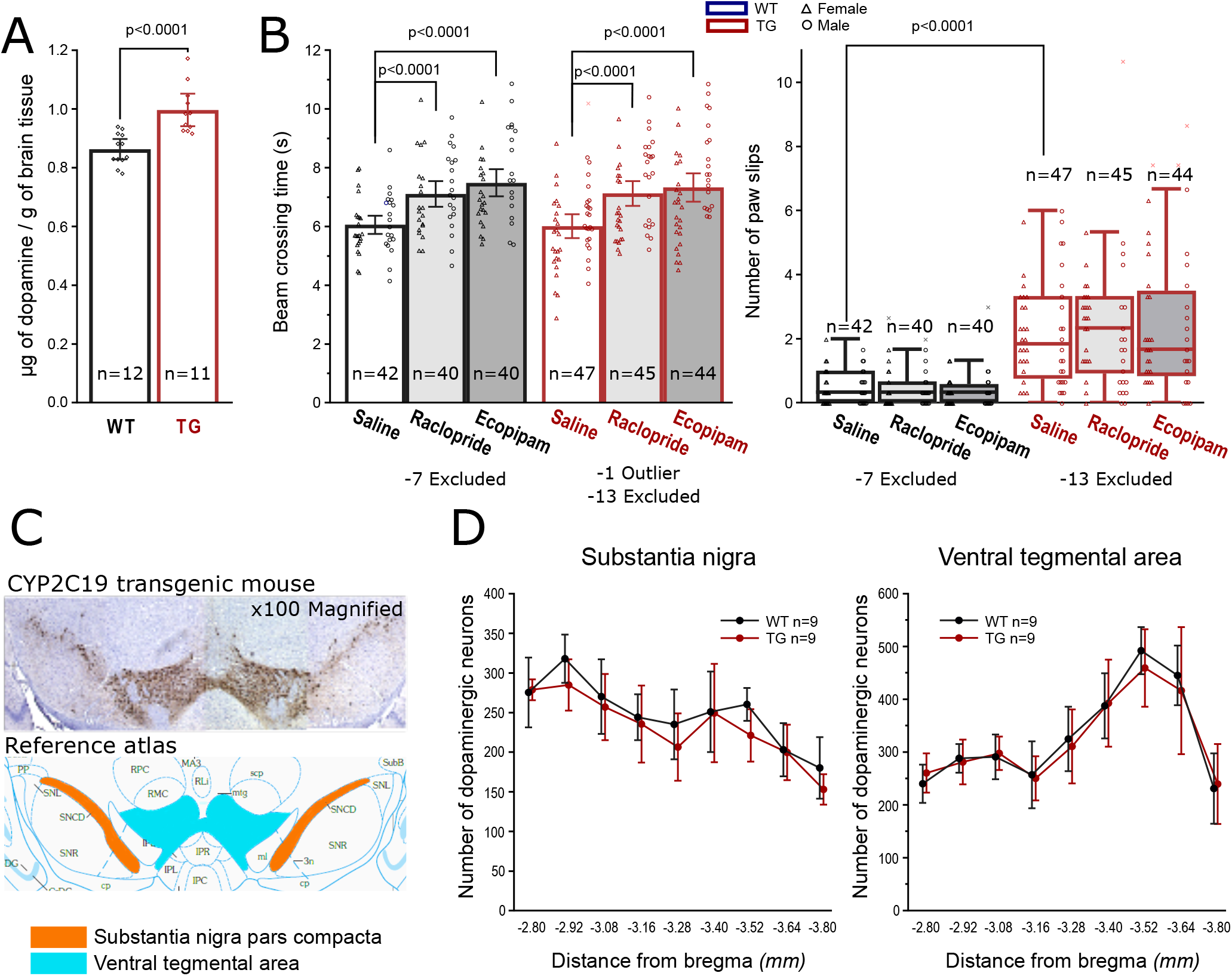
Subtle alterations of dopaminergic system in CYP2C19 transgenic mice (TG) compared to the wild type mice (WT). TG mice are hyperdopaminergic **(A)**, but dopaminergic antagonists failed to ameliorate their motoric impairment in the beam-walking test **(B).** Anti tyrosine hydroxylase (TH) immunohistochemistry **(C)** revealed no significant changes in the numbers of midbrain TH+ neurons **(D)** after FDR correction.

Next, potential neuroanatomical changes in TG mice were investigated throughout the brain by Gd-enhanced *ex-vivo* structural MRI neuroimaging. Significant, structural gray matter alterations identified through voxel-based morphometry (VBM) are graphically represented in 3D (Figure 3A) and as a collection of representative coronal neuroimaging slides (Figure 3B). Noteworthy however, the figures produced with VBM method represent total brain volume adjusted values. Since the total brain volume in the TG mice was significantly reduced by 3%, which is very substantial, volumes of the 20 segmented and analyzed regions of interests were not corrected for the total brain volume. Significant volumetric changes in TG compared to WT mice were observed in 10 out of 20 regions, out of which changes in 5 remained significant after multiple comparisons correction. Profound decrease (−11.8% [95%CI: −14.7%, −9.0%], *p*<0.0001, *q*<0.0001) was observed in TG cerebellar volume, while the TG hippocampal volume was moderately (−4.2% [95%CI: −6.4%, −1.9%], *p*=0.0004, *q*=0.015) reduced, compared to the corresponding volumes measured in WT mice. Also, moderate volumetric decreases in pons (−3.9% [95%CI: −5.6%, −2.1%], *p*<0.0001, *q*=0.0017) epithalamus (−4.6% [95%CI: −7.4%, −1.9%], *p*=0.0008, *q*=0.030) and inferior colliculus (−4.2% [95%CI: −6.4%, −1.9%], *p*=0.0004, *q*=0.015) were observed in TG compared to WT mice (Figure 3C). Effects of covariates on measured outcomes are presented in detail in the supplementary material. In summary, TG mice exhibit structural brain phenotype, which includes cerebellar and hippocampal volumetric decrease.

**Figure 3:**
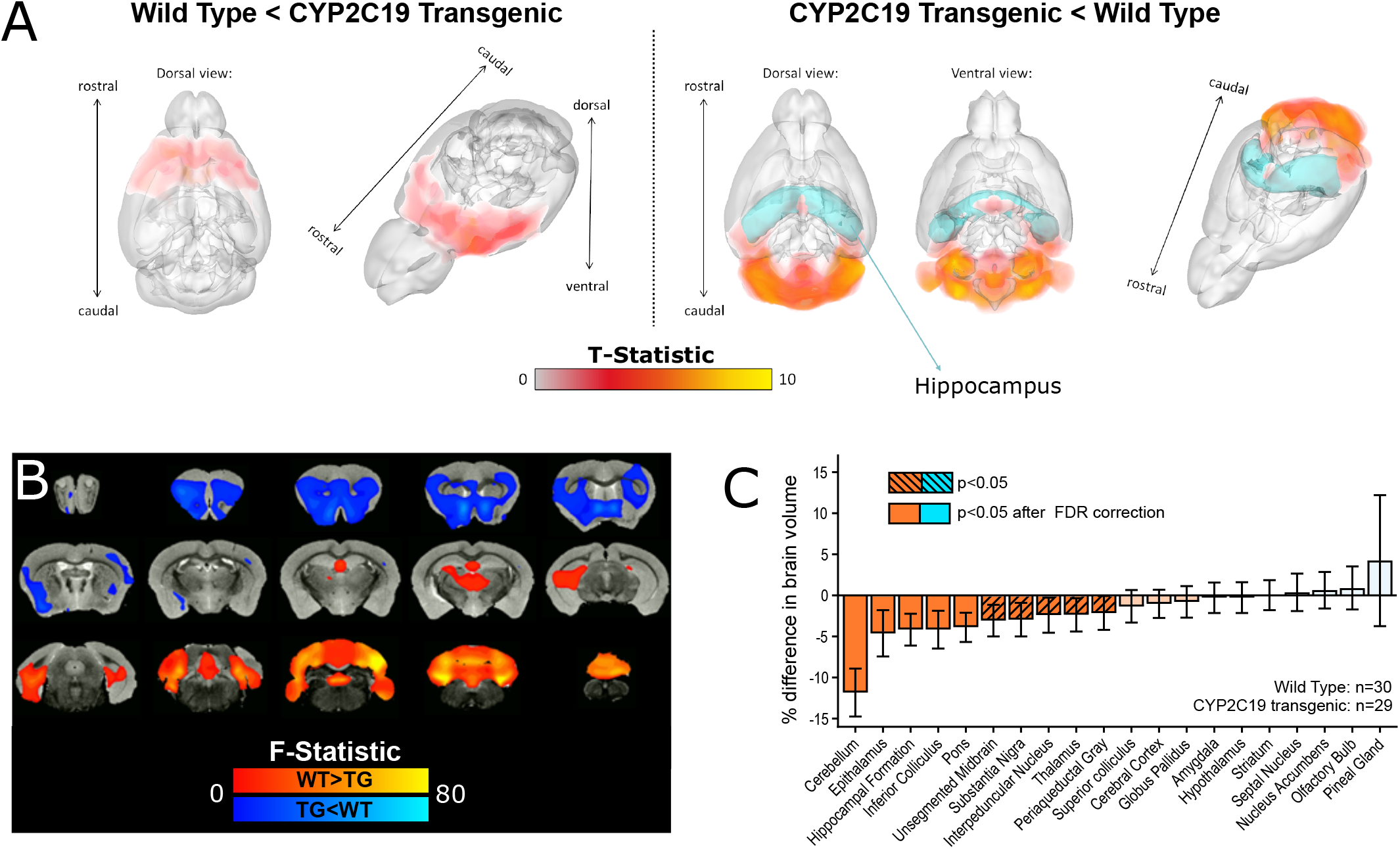
Structural changes in brains of 29 *CYP2C19* transgenic mice compared with 30 wild type mice measured with neuroimaging. Volumetric differences between *CYP2C19* transgenic mice and wild-types analyzed with voxel-based morphometry are presented as **(A)** 3D rendering and **(B)** coronal cross-sections. Panel **(C)** presents magnitudes of unadjusted volumetric changes in *CYP2C19* transgenic mice in 20 segmented regions of interest.

Finally, potential changes in antioxidative enzyme status within cerebellum and hippocampus were investigated. To indirectly investigate the presence of oxidative stress in cerebellum and hippocampus in TG mice, expression and activity of antioxidative enzymes: superoxide dismutase (SOD), catalase, glutathione peroxidase (GPx) and glutathione (GR) reductase in the tissues were measured. Activity of SOD was slightly increased (1.14-fold [CI95%: 1.06, 1.23], *p*=0.0010, *q*=0.023) in cerebellum and moderately increased (1.3-fold, [CI95%: 1.18, 1.47], *p*<0.0001, *q*=0.0013) in hippocampus of transgenic mice as compared to the controls. Furthermore, there was a significant increase (1.2-fold [CI95%: 1.13, 1.35], *p*<0.0001, *q*=0.0021) in GR enzyme activity in hippocampus of transgenic mice. On the other hand, there were no significant changes in the expression levels of any enzyme in the investigated brain regions measured with Western-blot method. In summary, TG mice exhibit normal expression but higher activity of anti-oxidative enzymes in cerebellum and hippocampus compared to the wild types. Detailed analysis outputs are presented in the Supplementary material.

## DISCUSSION

*CYP2C19* transgenic mouse (TG mouse) exhibits altered gait and slightly worse performance in more demanding motoric tasks, including impaired balance. This motoric phenotype shares several attributes of ataxia arguing for the face validity of the model; since abnormal walk, diminished balance and loss of fine dexterity are one of the most common symptoms of cerebellar ataxia^1^. Hyperdopaminergism observed in the TG mice is most likely not the cause of observed motoric genotype, but rather a compensatory response, since antidopaminergic drugs failed to ameliorate motoric impairments of TG mice and as hyperdopaminegism was not very profound in magnitude (Figure 2). Moreover, hyperdopaminergism was not associated with dopaminergic cell loss in the midbrain in both adult and older TG mice, despite hypothetically increased vulnerability of dopaminergic neurons to the chronic hyper-activation^20^. Even though neuromelanin aggregates are not normally detectable in rodents due to their relatively short lifespans compared to the long process of neuromelanin production^21^; few animal models with altered dopaminergic nigro-striatal pathway managed to exhibit detectable neuromelanin levels^21,22^. However, neuromelanin was not detected in 15-month TG mice in this study, arguing that the increase in dopamine production was not sufficient to cause any quantifiable neuromelanin aggregation.

Cerebellar atrophy observed in the TG mice is the most probable cause of ataxia-like phenotype, which is in concordance with neuroimaging observations in humans with ataxia^1^. Even though results in this report confirmed previous neuroimaging findings^6^ on reduced hippocampal volume in TG mice, Persson et. al. reported magnitude of 7.1% reduction in contrast to the 4.2% reduction observed here. This discrepancy most probably resulted from differences in tested cohorts; Persson et. al. included 7 male animals per group, while this study included 15 male and 15 female animals per group. The observed volumetric alterations may originate from aberrant development or may occur due to the proneness to atrophy of the affected structures after their development is completed. Increased activities of antioxidant enzymes in cerebellum and hippocampus of adult TG mice in our study (Figure 4) hints at increased oxidative stress and potential proneness to neuronal apoptosis in these regions.

**Figure 4:**
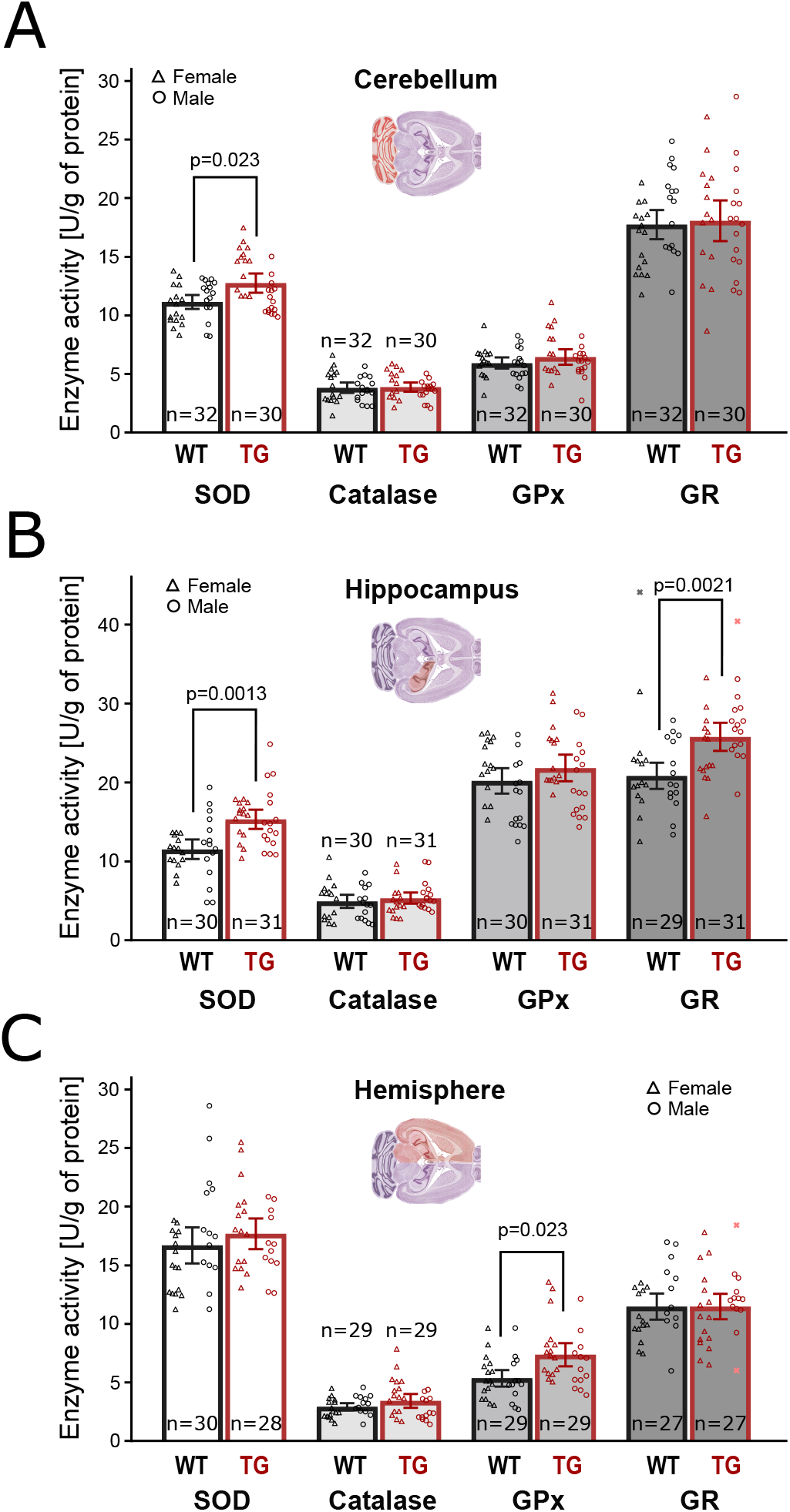
Antioxidant enzymes activity in 3 brain regions. Compared to controls (WT), *CYP2C19* transgenic mice (TG) exhibited **(A)** increased superoxide dismutase (SOD) activity in the cerebellum, **(B)** increased hippocampal activity of SOD and glutathione reductase (GR), and **(C)** increased activity of glutathione peroxidase (GPx) in the whole hemisphere.

Several animal models exhibit motoric phenotype similar to one observed in TG mice^23–25^. For example, Atcay^ji-hes^ mouse exhibits more extreme gait disturbances and involuntary hind-paw movements while walking, which were not observed in TG mice (Movie 2 from Luna-Cancalon et. al. 2014 ^23^). Next, *Reeler* mouse can be considered as the closest phenocopy of the TG mouse as it has the very similar gait (Video 1 in Machado et. al. 2020^24^); this behavior is also accompanied with impaired performance in rotarod and beam walking tests. *Reeler* mice also exhibit an aberrant formation of the hippocampus and severely reduced cerebellar size^24^. Finally, besides similar gait to TG mice, Sgce^m+/pGt^ mice also exhibits slight tremor, more slips in the beam walking test and abnormal expression of several genes in the cerebellum that are potentially associated with aberrant cerebellar development^25^. Considering the reported characteristics of phenotypically similar animal models, possible sources for gait abnormalities in TG mouse could lie in abnormal cerebellar development and/or reduced output from Purkinje cells, and further investigation of these functions in TG mice is needed. Importantly, the TG mice possess a few unique properties, not previously observed in other genetic models: **(1)** TG mouse loses the intensity of its motoric phenotype during maturation, while in most other animal models of ataxia nor progressive improvement nor constant levels of motoric impairments are apparent^4^; **(2)** While most animal models of ataxia exhibit altered movements in the both sides of the body^4^, in the TG mouse only one side of the body is affected with motoric impairment in the adulthood. In conclusion, CYP2C19 transgenic mice show ataxia-like phenotype and could be used in the preclinical research of new candidate-drugs for ataxia and cerebellar function and development in general. Understanding of the exact molecular mechanisms involved in TG mouse motoric phenotype, chiefly the processes behind the spontaneous improvement in the motoric phenotype in young TG mice, could provide insight into new drug targets for the treatment of cerebellar disorders.

### Limitations

Most importantly, the time course for several observed effects is still virtually unknown; measurement of brain volumes in different time points would be necessary in order to determine if the changes occurred during development or after birth and if this is the case, when exactly. Also, determination of dopamine levels in TG mice of different age would provide the time of onset and duration of hyperdopaminergism, while measurement of dopamine levels in the different brain areas would help to locate the sites of increased dopamine production. Next, antioxidative enzyme activity and expression only provides the information related to the general state in the observed regions; detection of ROS or oxidative damage is still needed to confirm presence of disrupted oxidative/antioxidative balance in brain regions of interest. In addition, systematic administration of antidopaminergic drugs leaves more room for confounding effects due to potency of these drugs and it is less informative compared to more precise drug administration techniques, such as stereotaxic injection into the dorsal striatum.

## Acknowledgement

We thank Nikola Ašujiċ and Teodora Miloševski for their contribution in conducting animal experiments.

## Funding sources for the study

This project received funding from The Ministry of education, science and technological development of the Republic of Serbia (Grant No 175036) to MMJ and VP; the Swedish Brain Foundation (Grant No FO2017-0111) to MIS and MMJ, and H2020 ERA-NET NEURON BrainCYP grant to MIS and RFT

https://farmacija-mv.sharepoint.com/personal/fmilosavljevic_pharmacy_bg_ac_rs/_layouts/15/onedrive.aspx?id=%2Fpersonal%2Ffmilosavljevic%5Fpharmacy%5Fbg%5Fac%5Frs%2FDocuments%2FAttachments%2FCYP%20Cerebellum%20%2D%20Gait%20video%281%29%2Emp4&parent=%2Fpersonal%2Ffmilosavljevic%5Fpharmacy%5Fbg%5Fac%5Frs%2FDocuments%2FAttachments

**Video 1: Ataxia-like gait in *CYP2C19* transgenic mouse**

Wild-type mouse was used as a control for the comparison of the gait disturbance in an adult (>10 weeks old) *CYP2C19* transgenic mouse, and more pronounced disturbance in 3-week old transgenic mouse.

https://farmacija-my.sharepoint.com/personal/fmilosavljevic_pharmacy_bg_ac_rs/_layouts/15/onedrive.aspx?id=%2Fpersonal%2Ffmilosavljevic%5Fpharmacy%5Fbg%5Fac%5Frs%2FDocuments%2FAttachments%2FCYP%20Cerebellum%20%2D%20Beam%20walking%20video%281%29%2Emp4&parent=%2Fpersonal%2Ffmilosavljevic%5Fpharmacy%5Fbg%5Fac%5Frs%2FDocuments%2FAttachments

**Video 2: *CYP2C19* transgenic mouse exhibits significantly more slips in the beam walking test**

Representative baseline beam-walking runs of one wild type and one *CYP2C19* transgenic mouse are shown. Criterion for the counting of the paw slips is also demonstrated.

## Notes

**Conflict of interest:** Authors report no conflict of interests

### Competing Interest Statement

The authors have declared no competing interest.

