## Supplementary Information for "Humanized *CYP2C19* transgenic mouse as an animal model of cerebellar ataxia"

### 1. METHODS

#### 1.1. Laboratory animals:

Animal wellbeing was checked minimum twice a week, but during the pharmacological treatments wellbeing and signs of distress were checked every day. Behavioral tests were performed during the light-phase and animals were accustomed to the testing room for at least 30 minutes. Diluted (10% v/v) ethanol was used to clean apparatus between the runs to eliminate the smell of a previous tested animal that could interfere with the test, while avoiding behavioral effects of inhaled ethanol.

#### 1.2. Clasping reflex screening:

Flexion of animal’s limbs or body or clinching of an animal’s paws together were used as visual signs of the clasping reflex, and both were induced once an animal was suspended by its tail and then slowly lowered towards the hard surface. On the other hand, extension of limbs in the anticipation of the fall was considered a physiological reaction^1^.

#### 1.3. Rotarod test protocol:

In the rotarod test, Ugo Basile Rotarod 47700 (Ugo Basile, Italy) apparatus with four 8.7cm wide lanes was used. Animals were trained to run against the rotating rod where the rotation speed was gradually increased until it became impossible for any mouse to keep the balance and stay on the rod. Two training trials were done on the two consecutive days with 3 runs per trial, and the animals that were unable to learn the task were excluded from the experiment. Every training run consisted of a 2-minute walk on a rod rotating at low speed of 5 RPM (Revolutions per Minute) with 5 minute breaks in the home cage between the runs. On a third day, animals were placed on the rod rotating at 2 RPM and were allowed to take a proper position, after which the test run was started and the rod began to accelerate at the rate of +8RPM per minute (2-50 RPM in 6 minutes). Falls were detected automatically by trip-switches under the rod; but if an animal grabbed onto the rod to avoid falling, the run was manually stopped at the moment an animal stopped running. Six runs were performed with at least 5 minute break in a home cage between the runs. Run was considered failed if an animal fell from the apparatus for any other reason besides failing to keep up with the rotation speed. If an animal had 3 or more failed runs, it was excluded from the experiment.

#### 1.4. Beam-walking test protocol:

In the beam-walking (BW) test, rectangular, 8mm wide and 1m long transparent acrylic beam was used. Strong light source was placed at the starting position on the beam in order to induce the anxious response in tested mice, and on the opposite end there was closed dark box which served as a goal, since mice have a natural tendency for seeking shelter in closed and dark spaces. Beam was also set at the slight incline since mice tend to seek high ground when escaping from a danger^2^. Mice were also allowed to accommodate to the dark box for 1 minute before every run in order to facilitate the feeling of safety inside the dark box. Animals were also trained to cross the beam for 2 consecutive days before the experiment. On the day 1, mice were placed 25cm, 50cm and 75cm away from the goal in the first, second and third training run, respectively; and in every run, mice were allowed to explore and find the way to the goal box. On the day 2, mice were allowed to cross the full length of the beam, and if an animal could not finish the task 3 times it was excluded from the experiment. On the third day, 5 test runs were performed with the additional sixth run, if at least one run failed. Run was considered failed if an animal slowed down or completely stopped while crossing the beam; if there were less than 3 valid runs in total animal was excluded from the experiment due to the lack of motivation to perform in this task. Every run was filmed for later analysis and beam crossing time was measured live using stopwatch, and validated later upon the analysis of the videos. Two researchers independently determined number of slips by reviewing the video footage, and one slip was defined as the failure to keep the paw on the beam while trying to make one step.

#### 1.5. Measurement of Whole-brain Dopamine Concentration:

Concentration of dopamine was determined in the brain hemispheres of 3 months old mice using HPLC-MS-MS method. Extraction solution (15% methanol and 15% acetonitrile in water adjusted to pH=2.5 using acetic acid) was added to the harvested mouse brains in the volume of 0.009ml/mg of tissue followed by homogenization. Dopamine-d4 was added to the homogenate as an internal standard in the amount of 1ng/mg of tissue followed by centrifugation at 11,000g. Twenty-five µl of supernatant was injected into the Agilent 1260 LC system containing (1) Agilent 1260 Quaternary pump set to flow rate of 0.200 ml/minute, (2) Agilent 1260 Infinity Standard Autosampler and (3) temperature-controlled column compartment that was connected to (4) Agilent 6430 QQQ. Gemini 5 µ C18 110 Å 150 x 2 mm column (Phenomenex, USA) was used and the mobile phase consisted of Solvent A (aqueous solution of acetic acid, pH 2.5) and Solvent B (methanol). The following gradient elution was used: 5% B 0 - 3 minutes, 3 - 5 minutes linear increase to 70% B, maintained 5-15 minutes, 15 - 16 minutes linear decrease to 5% B, maintained 16- 25 minutes. The QQQ was equipped with an electrospray ion source and operated in positive ion mode for the purpose of measuring dopamine with monitored transitions being 154 -> 137 for Dopamine and 158 -> 141 for Dopamine-d4. The calibration standards prepared in the extraction solution had concentration range of 5 - 1000 ng/ml^3^.

#### 1.6. Anti-dopaminergic Drugs and Motoric Performance

BW test was repeated after the treatment with selective D1 agonist ecopipam (SCH-39166, Tocris Bioscience, UK) and D2 antagonist raclopride (Tocris Bioscience, UK). Animals in the both test groups were divided into three subroups in a manner to ensure that the beam crossing time is equivalent between the subgroups. Each subgroup received saline, raclopride, or ecopipam prior to the retesting. There were three BW test trials with 7-day washout period between them, so that each animal could receive every respective treatment, as shown in the Table:

##### **Table S1:** Protocol of pharmacological treatment in the BW test

| Test groups (females/males) | Trial 1 |  | Trial 2 |  | Trial 3 |
| --- | --- | --- | --- | --- | --- |
| Group 1 n=30  CYP2C19 TG n=17 (9/8)  Wild-Type n=13 (7/6) | Saline | 7-day washout | Raclopride | 7-day washout | Ecopipam |
| Group 2 n=30  CYP2C19TG n=16 (8/8)  Wild-Type n=14 (8/6) | Raclopride |  | Ecopipam |  | Saline |
| Group 3: n=29  CYP2C19 TG n=15 (8/7)  Wild-Type n=14 (7/7) | Ecopipam |  | Saline |  | Raclopride |

The same BW test protocol was used as previously described, but without the training phase, since the animals already had previous experience with the BW test.

#### 1.7. Immunohistochemistry protocol:

Adult mouse brain samples from transgenic and control mice were fixed using 4% formaldehyde and embedded into paraffin blocks. Tissue samples were then cut into 10µm thick coronal sections using Leica RM 2125RT microtome (Leica Biosystems, Germany) that were collected on the Superfrost® Plus microscope slides (Thermo-Fisher Scientific, Germany). Initially, slides were deparaffinized by xylene and then rehydrated throughout the series of ethanol solutions of decreasing concentrations, followed by a brief rinse in the distilled water. From this point on, slides were rinsed with the PBS (Phosphate-buffered saline, 0.1M, pH=7.4) solution 3 times for 5 minutes between steps. Antigen retrieval was done by heating slides submerged into citrate buffer to its boiling point in the microwave oven for 20 minutes. To reduce unspecific staining due to endogenous peroxidase activity, slides were incubated with the 2% H2O2 solution for 10 minutes. Next, the slides were covered with blocking solution (2.5% Normal Horse Plasma, Vector Laboratories, USA) for 30 minutes to block potential signalling from nonspecific tissue epitopes. Next, slides were incubated overnight at +4°C with rabbit anti-Tyrosine Hydroxylase antibody (1:250 in PBS, ab137869, Abcam, UK). On the next day, slides were incubated with secondary horseradish peroxidase conjugated antibody (ImmPRESS™ Anti Rabbit Ig Reagent, Vector Laboratories, USA) for 2h. Specific signal was developed by using peroxidase substrate Diaminobenzidine (DAB substrate kit, ab64238, Abcam, UK) followed by brief counterstaining with haematoxylin stain. Representative slides were -2.80mm, -2.92mm, -3.08mm, -3.16mm, -3.28mm, -3.40mm, -3.52mm, -3.64mm and - 3.80mm from bregma (Figures 54, 55, 56, 57, 58, 59, 60, 61 and 62, respectively, from mouse brain atlas^4^. Stained slides were observed under the BX50 light microscope (Olympus System, Japan) with 3CCD Color Video Camera DXC-950P (Sony, Japan) mounted on the top. Composite micrographs were compared to the referent atlas in order to determine the boundaries of SN and VTA on the given section [Atlas].

#### 1.8. Neuromelanin detection protocol:

Presence of neuromelanin was assessed on 10µm thick sections of paraffin embedded mid-brains of 15-month-old transgenic and control mice stained with Fontana-Masson melanin stain (ab150669, Abcam, UK). In short, paraffinized sections were rehydrated and incubated at 60°C for about 40 minutes in freshly prepared ammoniac silver nitrate solution. Next, slides were incubated first with 0.2% gold-chloride solution for 30 seconds, then with sodium-thiosulfate for 2 minutes, and finally with nuclear Fast Red stain for 5 minutes at room temperature with rinsing in distilled water before every step. Stained slides were then dehydrated, mounted and observed under the BX50 light microscope (Olympus System, Japan). Presence of black granules in neurons was considered as a sign of neuromelanin aggregation.

#### 1.9. Gadolinium-enhanced Neuroimaging:

Gd containing contrast agent ProHance (0.5M Gadoteridol, Bracco diagnostics, USA) was used to increase the contrast during the MRI scans and consequently increase the resolution of the obtained 3D images. To prepare the brain tissue for neuroimaging animals were first overdosed with anaesthetic, fixed on the operation table, and thoracic cavity was opened in order to expose animal’s heart. Needle connected to the tubing of the peristaltic pump (LabKemi, Sweden) was inserted into left ventricle, and the cut was made on the right ventricle for the fluid outflow. Using the pump, whole animal was transcardially perfused first with 5ml of PBS solution in order to replace the blood and prevent clothing, followed with 10ml of Gd-enriched fixating solution (11.1% v/v of 36% formaldehyde solution and 10% v/v of contrast agent in 0.1M phosphate buffer solution, pH=7.4) which fixed the brain tissue and saturated brain extracellular fluid with Gd contrast agent. At the end of the perfusion, animals were decapitated and soft tissue, mandible and snout were removed, so that the skull with the intact brain inside remained as the final sample for neuroimaging. Harvested samples were kept in Gd enriched fixating solution overnight, followed by the transfer into gadolinium enriched PBS (1% v/v of ProHence in 100ml of 0.1M PBS, pH=7.4) in which samples were kept at +4°C until the MRI scan.

Medulla oblongata was also excluded due to possible damage to the tissue caused by decapitation of an animal. Volumes were analyzed for the following 20 regions of interest: Amygdala, Cerebellum, Cerebral Cortex, Corpus Striatum, Epithalamus, Globus Pallidus, Hippocampal Formation, Hypothalamus, Inferior Colliculus, Interpeduncular Nucleus, Nucleus Accumbens, Olfactory Bulb, Periaqueductal Gray, Pineal Gland, Pons, Septal Nucleus, Substantia Nigra, Superior Colliculus, Thalamus, Unsegmented Midbrain.

As of voxel-based morphometry, we had to generate a new study-specific mouse GM template, which was obtained by non-linearly registering all GM segmentations to the segmentation of one mouse, randomly chosen from the dataset, and averaging them. Afterwards, the original VBM pipeline was closely followed by registering all native GM images to the study-specific template and “modulating” them to correct for local expansion or contraction. The modulated GM images were then smoothed with an isotropic Gaussian kernel with a sigma of 0.3 mm.

#### 1.10. Antioxidant Enzyme Activity

Tris buffer (50 mM Tris, 0.25 M sucrose, 1 mM EDTA, pH 7.4) has been added to the flash frozen tissue in the volume of 0.1 ml/g of tissue prior to the homogenization by sonification (3 x 10s at 10 MHz on ice). This mixture was centrifuged at 105,000 g for 60 min and collected supernatant was stored in liquid nitrogen until use. The protein concentration was determined by the Lowry method^5^, and bovine serum albumin was used as standard. Total SOD activity was determined by the adrenaline method^6^ and SOD unit was defined as the catalytic activity that decreases the rate of adrenalin auto-oxidation by 50% at pH=10.2. CAT activity was determined according to Beutler^7^ and CAT unit was defined as the activity sufficient to decompose 1 mmol H2O2 per minute at 25°C and pH 7.0. The activity of GPx was determined by the glutathione reduction of t-butyl hydroperoxide, using the modification of the assay described by Paglia and Valenine^8^ and GR activity was determined using the method of Glatzle^9^. GPx unit was defined as the activity that oxidizes 1 mmol of NADPH per minute at 25°C and pH 7.0, and GR unit was defined as an enzyme activity that oxidizes 1nmol of NADPH per minute at 25°C and pH 7.4.

#### 1.11. Antioxidant Enzyme Expression

Western blot method for specific protein quantification was done by boiling of tissue samples in Laemmli's sample buffer, followed by protein resolution on the 10% or 12% SDS polyacrylamide gels. Western transfer of proteins from the gels to PVDF membranes was performed at 135 mA overnight in the 25 mM Tris buffer, pH 8.3 containing 192 mM glycine and 20% v/v methanol. The membanes were blocked with the 5% non-fat dry milk in PBS (1.5 mM KH2PO4, 6.5 mM Na2HPO4, pH 7.2, 2.7 mM KCl, 0.14 M NaCl) for 1h at room temperature. After blocking, membranes were incubated overnight at 4 °C with the respective primary antibody: SOD1 (ab13498, 1:5000), SOD2 (ab13533, 1:2000), CAT (ab16731, 1:2000), GPx (ab22604, 1:2000), GR (ab16801, 1:2000), and β actin (ab8227, 1:5000). Membranes were subsequently washed with PBS containing 0.1% Tween 20 and incubated for 1h at room temperature with goat anti-rabbit horseradish peroxidase-conjugated secondary antibody (ab6721, 1:30000). All antibodies were purchased from Abcam, UK. The immunoreactive proteins were visualized by chemiluminiscent method using iBright Western Blot Imaging Systems (Thermo-Fisher Scientific, Germany). Quantitative analysis was performed using iBright Analysis Software (Thermo-Fisher Scientific, Germany). β-actin was used as equal load control.

#### 1.12. Statistics:

To investigate the genotype-specific effects on experimental readouts, Student’s t-test for independent samples was used. If experiment included a covariate, one-way ANCOVA test was used with genotype status as independent fixed variable, and sex or age as covariate; if a covariate did not significantly affect the results; it was dropped from the analysis. For the experiments that included drug treatment, a two-way mixed ANCOVA was used with the treatment as the repeated-measurement independent variable, genotype as the fixed independent variable, and sex as the covariate. In the case of non-normally distributed data, non-parametric alternatives (Mann Witney and Kuskall Wallis tests) were used.

### 2. RESULTS:

#### 2.1. Tabular representation of genotype specific results of the study

##### **Table S2:** Significance and magnitude of the impact of the genotype on the study results:

| Test | Readout | Number of animals  (Outliers; Excluded) | | | %(2C19TG-WT)  [CI95%] | Sign. level |
| --- | --- | --- | --- | --- | --- | --- |
| Rotarod | Latency to fall | | n=97  (O=2) | +0.5%  [-7.1%, +8.2%] | | p=0.90 |
| Beam walking | Beam-crossing time | | n=85  (O=4; E=5*) | +14%  [+6.4%, +22%] | | p=0.0014 |
|  | Number of slips | | n=89  (E=5*) | +457% | | p<0.0001 |
| Beam walking  i.p. Saline | Beam-crossing time | | n=89  (E=4*) | -2.2%  [-10%, +5.5%] | | p=0.81 |
|  | Number of slips | | n=89  (E=4*) | +440% | | p<0.0001 |
| Beam walking  i.p. Raclopride | Beam-crossing time | | n=85  (E=8*) | -0.5%  [-8.8%, +7.8%] | | p=0.81 |
|  | Number of slips | | n=85  (E=8*) | +590% | | p<0.0001 |
| Beam walking  i.p. Ecopipam | Beam-crossing time | | n=84  (E=9*) | -2.6%  [-11%, +6.1%] | | p=0.81 |
|  | Number of slips | | n=84  (E=9*) | +457% | | p<0.0001 |
| Dopamine levels | µl/g of tissue | | n=23 | +15%  [+12%, 19%] | | p<0.0001 |
| **Gd-enhanced neuroimaging** | | | | | | |
| Amygdala volume | Number of voxels | | n=59 | -0.3%  [-2.1, +1.6%] | | p=0.77  q=1.0 |
| Cerebellum volume | Number of voxels | | n=56  (O=3) | -12%  [-15%, -8.9%] | | p<0.0001  q<0.0001 |
| Cerebral Cortex volume | Number of voxels | | n=58  (O=1) | -1.0%  [-2.8%, +0.7%] | | p=0.22  q=1.0 |
| Corpus Striatum volume | Number of voxels | | n=59 | +0.0%  [-1.8%, +1.8%] | | p=0.99  q=1.0 |
| Epithalamus volume | Number of voxels | | n=59 | -4.6%  [-7.4%, -1.8%] | | p=0.0008  q=0.030 |
| Globus Pallidus volume | Number of voxels | | n=59 | -0.8%  [-2.7%, +1.1%] | | p=0.41  q=1.0 |
| Hippocampus volume | Number of voxels | | n=59 | -4.2%  [-6.1%, -2.2%] | | p<0.0001  q=0.0027 |

continued **Table S2:** Significance and magnitude of the impact of the genotype on the study results:

| Test | Readout | Number of animals  (Outliers; Excluded) | | | %(2C19TG-WT)  [CI95%] | Sign. level |
| --- | --- | --- | --- | --- | --- | --- |
| Hypothalamus volume | Number of voxels | | n=59 | -0.3%  [-2.2%, +1.6%] | | p=0.79  q=1.0 |
| Inferior Colliculus volume | Number of voxels | | n=59 | -4.2%  [-6.5%, -1.9%] | | p=0.0004  q=0.015 |
| Interpeduncular Nucleus volume | Number of voxels | | n=59 | -2.4%  [-4.5%, -0.3%] | | p=0.030  q=0.89 |
| Nucleus Accumbens volume | Number of voxels | | n=59 | +0.6%  [-1.6%, +2.8%] | | p=0.56  q=1.0 |
| Olfactory Bulb volume | Number of voxels | | n=58  (O=1) | 0.9%  [-1.7%, +3.5%] | | p=0.43  q=1.0 |
| Periaqueductal Gray Matter volume | Number of voxels | | n=54  (O=5) | -2.1%  [-4.2%, -0.1%] | | p=0.40  q=1.0 |
| Pineal Gland volume | Number of voxels | | n=59 | +4.2%  [-3.8%, +12%] | | p=0.29  q=1.0 |
| Pons volume | Number of voxels | | n=59 | -3.9%  [-5.7%, -2.1%] | | p<0.0001  q=0.0017 |
| Septal Nucleus volume | Number of voxels | | n=59 | +0.4%  [-1.9%, +2.7%] | | p=0.76  q=1.0 |
| Substantia Nigra volume | Number of voxels | | n=59 | -3.0%  [-5.0%, -0.9%] | | p=0.006  q=0.18 |
| Superior Colliculus volume | Number of voxels | | n=59 | -1.3%  [-3.3%, +0.6%] | | p=0.18  q=1.0 |
| Thalamus volume | Number of voxels | | n=59 | -2.4%  [-4.4%, -0.4%] | | p=0.022  q=0.68 |
| Unsegmented Midbrain volume | Number of voxels | | n=59 | -3.1%  [-5.0%, -1.2%] | | p=0.0023  q=0.077 |
| **Antioxidant enzyme activity** | | | | | | |
| SOD (hemisphere) | U/g of tissue | | n=58  (O=1) | +6.0%  [-5.6%, +18%] | | p=0.30  q=1.0 |
| SOD (cerebellum) | U/g of tissue | | n=62 | +14%  [+5.8%, +23%] | | p=0.0010  q=0.021 |
| SOD (hippocampus) | U/g of tissue | | n=61 | +33%  [+18%, +47%] | | p<0.0001  q=0.0013 |
| Catalase (hemisphere) | U/g of tissue | | n=58 | +17%  [-4.6%, +39%] | | p=0.12  q=1.0 |
| Catalase (cerebellum) | U/g of tissue | | n=62 | +1.4%  [-13%, +16%] | | p=0.43  q=1.0 |
| Catalase (hippocampus) | U/g of tissue | | n=61 | +8.8%  [-4.2%, +21%] | | p=0.82  q=1.0 |
| GPx (hemisphere) | U/g of tissue | | n=58 | +38%  [+16%, +59%] | | p=0.0011  q=0.023 |

continued **Table S2:** Significance and magnitude of the impact of the genotype on the study results:

| Test | Readout | Number of animals  (Outliers; Excluded) | | | %(2C19TG-WT)  [CI95%] | Sign. level |
| --- | --- | --- | --- | --- | --- | --- |
| GPx (cerebellum) | U/g of tissue | | n=62 | +8.8%  [-13%, +30%] | | p=0.16  q=1.0 |
| GPx (hippocampus) | U/g of tissue | | n=61 | +8.1%  [-2.9%, +19%] | | p=0.11  q=1.0 |
| GR (hemisphere) | U/g of tissue | | n=54  (O=3) | +0.0%  [-13%, +13%] | | p=0.91  q=1.0 |
| GR (cerebellum) | U/g of tissue | | n=63 | +1.8%  [-9.7%, +13%] | | p=0.78  q=1.0 |
| GR (hippocampus) | U/g of tissue | | n=60  (O=1) | +23%  [+13%, +35%] | | p<0.0001  q=0.0021 |
| **Antioxidant enzyme expression** | | | | | | |
| SOD1 (hemisphere) | Arbitrary Units | | n=48 | +12%  [-1.4%, +25%] | | p=0.084  q=1.0 |
| SOD1 (cerebellum) | Arbitrary Units | | n=41  (O=7) | +10%  [-5.3%, +26%] | | p=0.26  q=1.0 |
| SOD1 (hippocampus) | Arbitrary Units | | n=48 | +5.2%  [-11%, +16%] | | p=0.52  q=1.0 |
| SOD2 (hemisphere) | Arbitrary Units | | n=47  (O=1) | +5.7%  [-4.7%, +16%] | | p=0.28  q=1.0 |
| SOD2 (cerebellum) | Arbitrary Units | | n=44  (O=2, E=2^+^) | +23%  [+7.9%, +39%] | | p=0.0074  q=0.21 |
| SOD2 (hippocampus) | Arbitrary Units | | n=46  (O=2) | -4.3%  [-14%, +4.9%] | | p=0.36  q=1.0 |
| Catalase (hemisphere) | Arbitrary Units | | n=43  (O=5) | -0.2%  [-7.2%, +6.8%] | | p=0.98  q=1.0 |
| Catalase (cerebellum) | Arbitrary Units | | n=46  (E=2^+^) | +6.5%  [-6.2%, +19%] | | p=0.32  q=1.0 |
| Catalase (hippocampus) | Arbitrary Units | | n=35  (O=1, E=12^+^) | -0.1%  [-18%, +18%] | | p=0.98  q=0.98 |
| GPx (hemisphere) | Arbitrary Units | | n=48 | +3.8%  [-9.8%, +17%] | | p=0.58  q=1.0 |
| GPx (cerebellum) | Arbitrary Units | | n=46  (O=2) | -3.5%  [-12%, +5.2%] | | p=0.44  q=1.0 |
| GPx (hippocampus) | Arbitrary Units | | n=48 | +6.7%  [-5.2%, +19%] | | p=0.27  q=1.0 |
| GR (hemisphere) | Arbitrary Units | | n=48 | -5.4%  [-15%, +4.2%] | | p=0.28  q=1.0 |
| GR (cerebellum) | Arbitrary Units | | n=48 | +4.7%  [-9.7%, +16%] | | p=0.52  q=1.0 |
| GR (hippocampus) | Arbitrary Units | | n=47  (O=1) | +1.6%  [-9.4%, +13%] | | p=0.79  q=1.0 |

continued **Table S2:** Significance and magnitude of the impact of the genotype on the study results:

| Test | Readout | Number of animals  (Outliers; Excluded) | | | %(2C19TG-WT)  [CI95%] | Sign. level |
| --- | --- | --- | --- | --- | --- | --- |
| **Number of midbrain dopaminergic neurons** | | | | | | |
| Coronal section  -2.80mm from Bregma | Substantia Nigra | | n=18 | **+1.3%**  **[-12.9%, +15.5%]** | | **p=0.86**  **q=1.0** |
|  | Ventral Tegmental Area | | n=18 | **+8.3%**  **[-10%, +27%]** | | **p=0.36**  **q=1.0** |
| Coronal section  -2.92mm from Bregma | Substantia Nigra | | n=18 | **-10.5%**  **[-22.4%, +1.4%]** | | **p=0.11**  **q=0.80** |
|  | Ventral Tegmental Area | | n=18 | **-2.4%**  **[-17%, -13%]** | | **p=0.77**  **q=1.0** |
| Coronal section  -3.08mm from Bregma | Substantia Nigra | | n=18 | **-4.9%**  **[-24.7%, +14.9%]** | | **p=0.65 q=1.0** |
|  | Ventral Tegmental Area | | n=18 | **2.3%**  **[-13%, -18%]** | | **p=0.75**  **q=1.0** |
| Coronal section  -3.16mm from Bregma | Substantia Nigra | | n=18 | **-3.5%**  **[-23.3%, +16.4%]** | | **p=0.69**  **q=1.0** |
|  | Ventral Tegmental Area | | n=18 | **-2.6%**  **[-28%, +22%]** | | **p=0.83**  **q=1.0** |
| Coronal section  -3.28mm from Bregma | Substantia Nigra | | n=18 | **-12.2%**  **[-34.4%, +9.9%]** | | **p=0.19**  **q=1.0** |
|  | Ventral Tegmental Area | | n=18 | **-4.4%**  **[-29%, +20%]** | | **p=0.56**  **q=1.0** |
| Coronal section  -3.40mm from Bregma | Substantia Nigra | | n=18 | **-0.6%**  **[-27.8%, +26.6%]** | | **p=0.96**  **q=1.0** |
|  | Ventral Tegmental Area | | n=18 | **+1.3%**  **[%-21, +8.1%]** | | **p=0.92**  **q=1.0** |
| Coronal section  -3.52mm from Bregma | Substantia Nigra | | n=18 | **-15.0%**  **[-27.8%, -2.1%]** | | **p=0.037**  **q=0.18** |
|  | Ventral Tegmental Area | | n=18 | **-6.7%**  **[-21%, +8.1%]** | | **p=0.39**  **q=1.0** |
| Coronal section  -3.64mm from Bregma | Substantia Nigra | | n=18 | **-1.7%**  **[-22.1%, 18.6%]** | | **p=0.83**  **q=1.0** |
|  | Ventral Tegmental Area | | n=18 | **-6.4%**  **[-32%, +19%]** | | **p=0.63**  **q=1.0** |
| Coronal section  -3.80mm from Bregma | Substantia Nigra | | n=17  (O=1) | **-6.9%**  **[-20.1%, +6.3%]** | | **p=0.15**  **q=1.0** |
|  | Ventral Tegmental Area | | n=17  (O=1) | **+1.7%**  **[-14%, +18%]** | | **p=0.83**  **q=1.0** |

*Excluded due to failed test.

#### 2.2. Effects of covariates:

Statistical significance of the effect of covariates on the results is presented in the TableS2 and TableS3:

##### **Table S3:** Significance of the impact of sex as a covariate in the study results

| Test | *Readout* | Total number of animals | Significance |
| --- | --- | --- | --- |
| Rotarod | Latency to fall | n=97 | p=0.0001 |
| Beam walking | Beam-Crossing time | n=85 | p=0.70 |
|  | Number of slips |  | p=0.26 |
| Beam walking – i.p. Saline | Beam-Crossing time | n=90 | p=0.11 |
|  | Number of slips |  | p=0.79 |
| Beam walking – i.p. Raclopride | Beam-Crossing time | n=85 | p=0.030 |
|  | Number of slips |  | p=0.44 |
| Beam walking – i.p. Ecopipam | Beam-Crossing time | n=84 | p=0.0008 |
|  | Number of slips |  | p=0.58 |
| Dopamine levels | µl/g of tissue | n=23 | N/A |
| Gd-enhanced neuroimaging | | | |
| Amygdala volume | Number of voxels | n=59 | p=0.44  q=1.0 |
| Cerebellum volume | Number of voxels | n=56 | p=0.030  q=0.87 |
| Cerebral Cortex volume | Number of voxels | n=58 | p=0.070  q=1.0 |
| Corpus Striatum volume | Number of voxels | n=59 | p=0.27  q=1.0 |
| Epithalamus volume | Number of voxels | n=59 | p=0.010  q=0.33 |
| Globus Pallidus volume | Number of voxels | n=59 | p=0.72  q=1.0 |
| Hippocampus volume | Number of voxels | n=59 | p=0.58  q=1.00 |
| Hypothalamus volume | Number of voxels | n=59 | p=0.19  q=1.0 |
| Inferior Colliculus volume | Number of voxels | n=59 | p=0.047  q=1.0 |
| Interpeduncular Nucleus volume | Number of voxels | n=59 | p=0.41  q=1.0 |
| Nucleus Accumbens volume | Number of voxels | n=59 | p=0.35  q=1.0 |
| Olfactory Bulb volume | Number of voxels | n=58 | p=0.0022  q=0.078 |

continued **Table S3**: Significance of the impact of sex as a covariate in the study results

| Test | *Readout* | Total number of animals | Significance |
| --- | --- | --- | --- |
| Periaqueductal Gray Matter volume | Number of voxels | n=54 | p=0.51  q=1.0 |
| Pineal Gland volume | Number of voxels | n=59 | p=0.32  q=1.0 |
| Pons volume | Number of voxels | n=59 | p=0.13  q=1.0 |
| Septal Nucleus volume | Number of voxels | n=59 | p=0.82  q=1.0 |
| Substantia Nigra volume | Number of voxels | n=59 | p=0.41  q=1.0 |
| Superior Colliculus volume | Number of voxels | n=59 | p=0.57  q=1.0 |
| Thalamus volume | Number of voxels | n=59 | p=0.12  q=1.0 |
| Unsegmented Midbrain volume | Number of voxels | n=59 | p=0.70  q=1.0 |
| Antioxidant enzyme activity | | | |
| SOD (hemisphere) | U/g of tissue | n=58 | p=0.35  q=1.0 |
| SOD (cerebellum) | U/g of tissue | n=62 | p=0.030  q=0.58 |
| SOD (hippocampus) | U/g of tissue | n=61 | p=0.70  q=1.0 |
| Catalase (hemisphere) | U/g of tissue | n=58 | p=0.15  q=1.0 |
| Catalase (cerebellum) | U/g of tissue | n=62 | p=0.13  q=1.0 |
| Catalase (hippocampus) | U/g of tissue | n=61 | p=0.89  q=1.0 |
| GPx (hemisphere) | U/g of tissue | n=58 | p=0.18  q=1.0 |
| GPx (cerebellum) | U/g of tissue | n=62 | p=0.078  q=1.0 |
| GPx (hippocampus) | U/g of tissue | n=61 | p=0.0014  q=0.029 |
| GR (hemisphere) | U/g of tissue | n=54 | p=0.069  q=1.0 |
| GR (cerebellum) | U/g of tissue | n=63 | p=0.26  q=1.0 |
| GR (hippocampus) | U/g of tissue | n=60 | p=0.12  q=1.0 |

**continued Table S3:** Significance of the impact of sex as a covariate in the study results

| Test | *Readout* | Total number of animals | Significance |
| --- | --- | --- | --- |
| Antioxidative enzyme expression | | | |
| SOD1 (hemisphere) | Arbitrary Units | n=48 | p=0.23  q=1.0 |
| SOD1 (cerebellum) | Arbitrary Units | n=41 | p=0.45  q=1.0 |
| SOD1 (hippocampus) | Arbitrary Units | n=48 | p=0.31  q=1.0 |
| SOD2 (hemisphere) | Arbitrary Units | n=47 | p=0.0039  q=0.12 |
| SOD2 (cerebellum) | Arbitrary Units | n=44 | p=0.95  q=1.0 |
| SOD2 (hippocampus) | Arbitrary Units | n=46 | p=0.68  q=1.0 |
| Catalase (hemisphere) | Arbitrary Units | n=43 | p=0.59  q=1.0 |
| Catalase (cerebellum) | Arbitrary Units | n=46 | p=0.50  q=1.0 |
| Catalase (hippocampus) | Arbitrary Units | n=35 | p=0.49  q=1.0 |
| GPx (hemisphere) | Arbitrary Units | n=48 | p=0.82  q=1.0 |
| GPx (cerebellum) | Arbitrary Units | n=46 | p=0.78  q=1.0 |
| GPx (hippocampus) | Arbitrary Units | n=48 | p=0.18  q=1.0 |
| GR (hemisphere) | Arbitrary Units | n=48 | p=0.61  q=1.0 |
| GR (cerebellum) | Arbitrary Units | n=48 | p=0.67  q=1.0 |
| GR (hippocampus) | Arbitrary Units | n=47 | p=0.35  q=1.0 |

##### **Table S4:** Significance of the impact of age as a covariate in the study results

| Test | *Readout* | Total number of animals | Significance |
| --- | --- | --- | --- |
| Coronal section  -2.80mm from Bregma | Number of neurons in  Substantia Nigra | n=18 | p=0.58  q=1.0 |
|  | Number of neurons in  Ventral Tegmental Area | n=18 | p=0.079  q=0.32 |
| Coronal section  -2.92mm from Bregma | Number of neurons in  Substantia Nigra | n=18 | p=0.83  q=1.0 |
|  | Number of neurons in  Ventral Tegmental Area | n=18 | p=0.70  q=1.0 |
| Coronal section  -3.08mm from Bregma | Number of neurons in  Substantia Nigra | n=18 | p=0.94  q=1.0 |
|  | Number of neurons in  Ventral Tegmental Area | n=18 | p=0.044  q=0.13 |
| Coronal section  -3.16mm from Bregma | Number of neurons in  Substantia Nigra | n=18 | p=0.014  q=0.056 |
|  | Number of neurons in  Ventral Tegmental Area | n=18 | p=0.18  q=0.89 |
| Coronal section  -3.28mm from Bregma | Number of neurons in  Substantia Nigra | n=18 | p=0.005  q=0.005 |
|  | Number of neurons in  Ventral Tegmental Area | n=18 | p<0.0001  q<0.0001 |
| Coronal section  -3.40mm from Bregma | Number of neurons in  Substantia Nigra | n=18 | p=0.006  q=0.012 |
|  | Number of neurons in  Ventral Tegmental Area | n=18 | p=0.52  q=1.0 |
| Coronal section  -3.52mm from Bregma | Number of neurons in  Substantia Nigra | n=18 | p=0.33  q=1.0 |
|  | Number of neurons in  Ventral Tegmental Area | n=18 | p=0.28  q=1.0 |
| Coronal section  -3.64mm from Bregma | Number of neurons in  Substantia Nigra | n=18 | p=0.006  q=0.018 |
|  | Number of neurons in  Ventral Tegmental Area | n=18 | p=0.39  q=1.0 |
| Coronal section  -3.80mm from Bregma | Number of neurons in  Substantia Nigra | n=17 | p=0.11  q=0.65 |
|  | Number of neurons in  Ventral Tegmental Area | n=17 | p=0.040  q=0.080 |

*Excluded for lack of motivation to complete the task of the failure to complete the training,

^+^Excluded for the invalid measurement in the given sample
2.3. Determination of the cohort size

One mouse was considered as a single experimental unit. Three separate mice cohorts, each consisting out of one experimental (CYP2C19 transgenic mice - TG) and one control (wild type - WT) group, were used for this study:

- Cohort 1: For the whole-brain dopamine concentration measurement
  - n=23; 12 WT + 11 TG; all males; 3 months old mice
  - Procedure: Sacrificed for the sampling of the entire brain
- Cohort 2: For the immunohistochemistry and neuromelanin staining
  - n=18; 4 WT + 4 TG (15-month-old); 5 TG + 5 WT (6-month-old); all males
  - Procedure: Sacrificed for the production of the histological brain microscope slides
- Cohort 3: For the purpose of the motoric behaviour tests, neuroimaging, and antioxidative enzyme status
  - n= 99; 47WT + 52 TG; 3-6 months old; males+females
  - Procedure: First, rotarod test was performed followed with baseline beam-walking test.
  - Then, beam-walking test was repeated under antidopaminergic treatment protocol described previously.
  - Then, 60 out of 99 animals were sacrificed for the purpose of postmortem neuroimaging.
  - Remaining 39 animal cohort was expanded with 25 more animals (12 WT + 13 TG, 3-6 months old), and this groups of 64 animals was sacrificed for the sampling of the 3 brain regions for the measurement of the antioxidative enzyme expression and activity.

Number of animals for the cohort 3, used for the majority of the experiments, was decided based on the standard formula^10^:

n = $\frac{{(Z_{\alpha/2}+Z_{\beta})}^{2}\times2\times\sigma^{2}}{d^{2}}$

where the target difference between the groups (*d*) was set at 10%, since *CYP2C19* transgenic mice were expected to show mild motoric impairment of around 10% in the rotarod and beam-walking test, if any. Also, standard deviation (σ) of 20% was chosen for the calculations based on the result of the pilot study^11^, confidence level was set at 95% and power to 80%. This analysis predicted that 126 animals (63 TG + 63 WT) were needed for the experiment with the given parameters, but this number was limited to 100 animals due to practical reasons. Since large animal cohort was needed, special care was taken to re-use the same cohort in as many experiments in order to reduce animal suffering as much as possible.

Other cohorts were selected in a way to produce most meaningful results with as few animals as possible.

#### 2.4. Expression of the antioxidative enzymes in the brain tissue of CYP2C19 transgenic mice

##### **Figure S1:** Expression of antioxidant enzymes in 3 brain regions.

There was no change between CYP2C19 transgenic and Wild type mice in the exoression of either **(A)** Cupper-Zink Superoxide dismutase (SOD1), **(B)** Manganese Superoxide dismutase (SOD2), **(C)** Catalase **(D)** Glutathione peroxidase (GPx) and **(E)** Glutathione reductase (GR) in whole hemisphere, cerebellum or hippocampus. Primary analysis showed significant increase in the expression of SOD2 enzyme in cerebellum (23% increase CI95%: [7.9%, 39%], p=0.0074) in TG mice but this change did not remain significant after correction for multiple comparisons (q=0.12). Enzyme relative concentrations were determined by western blot method and concentrations were expressed in arbitrary units.

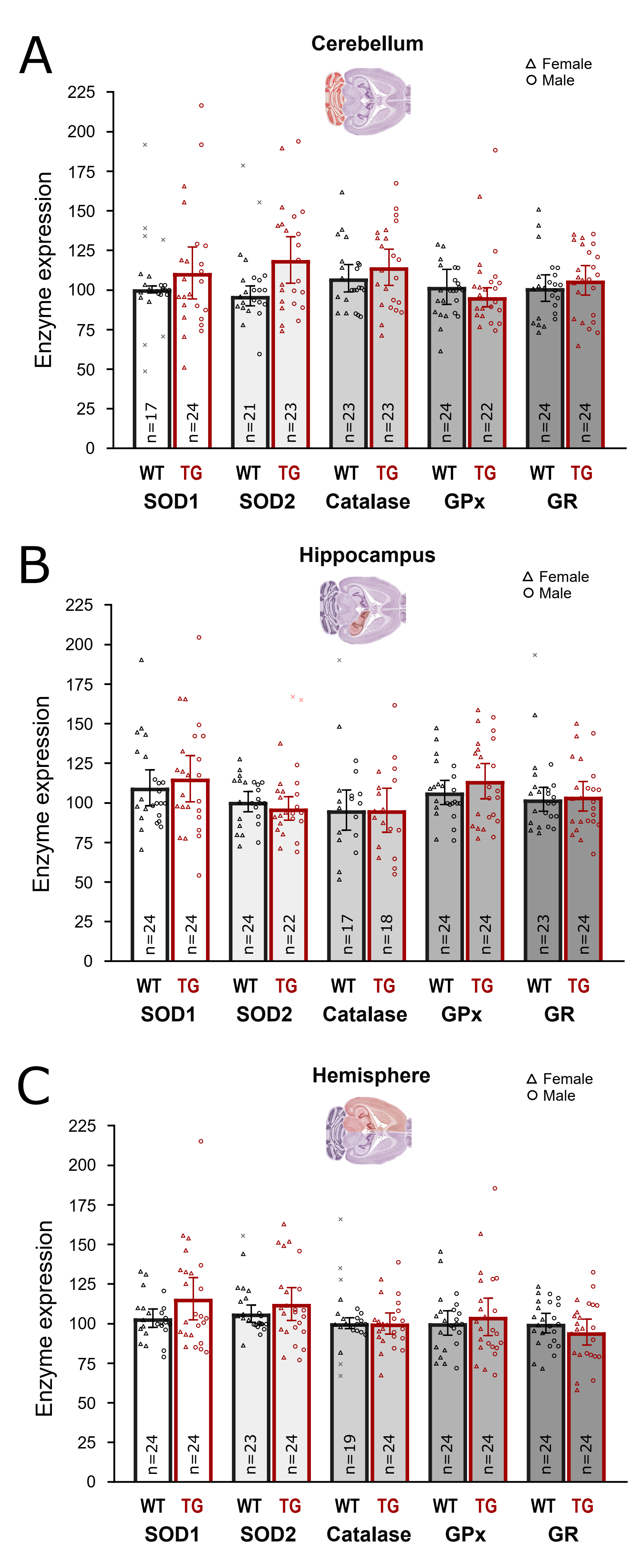

#### 2.5. ARRIVE 2.0 guidelines checklist

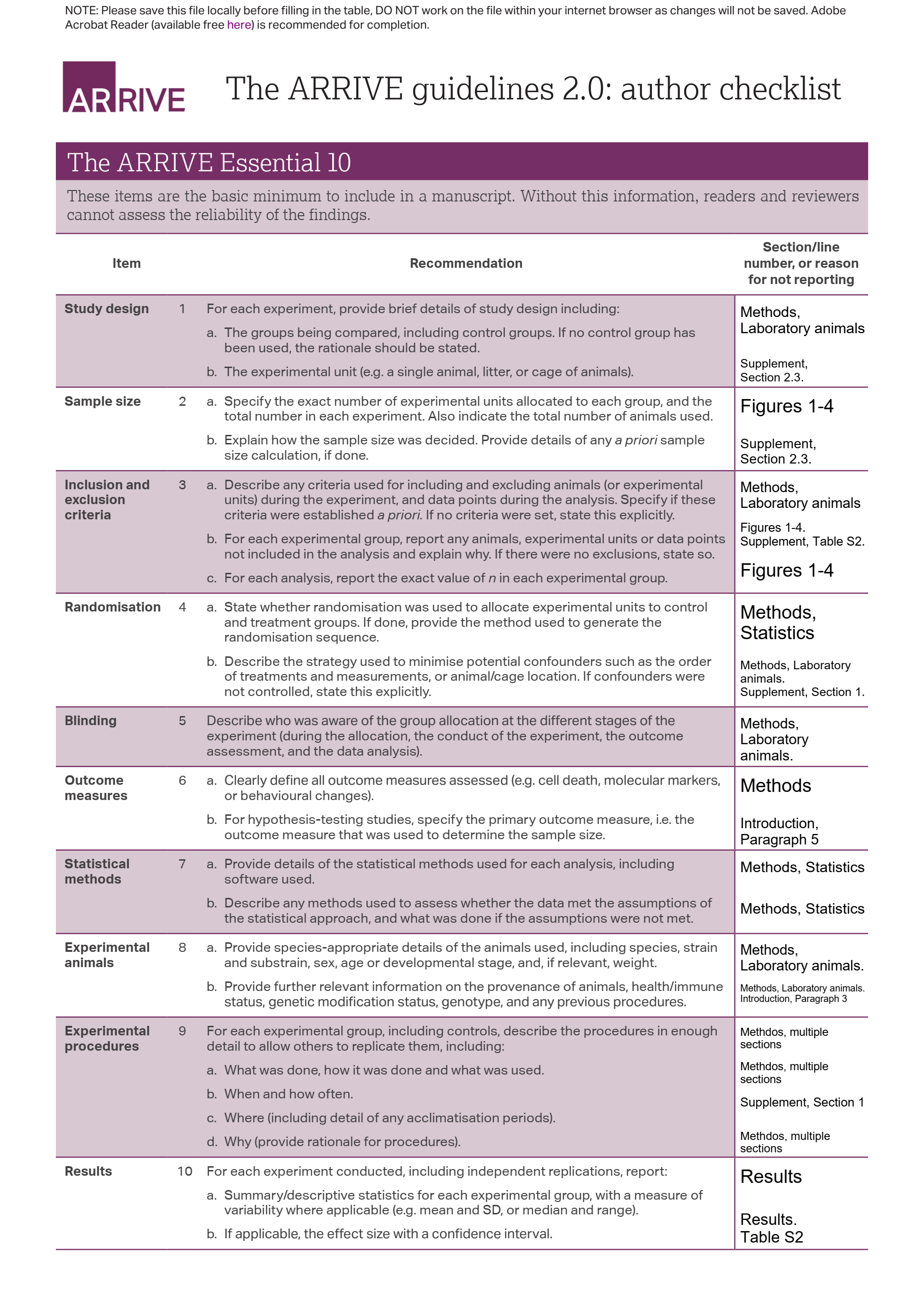

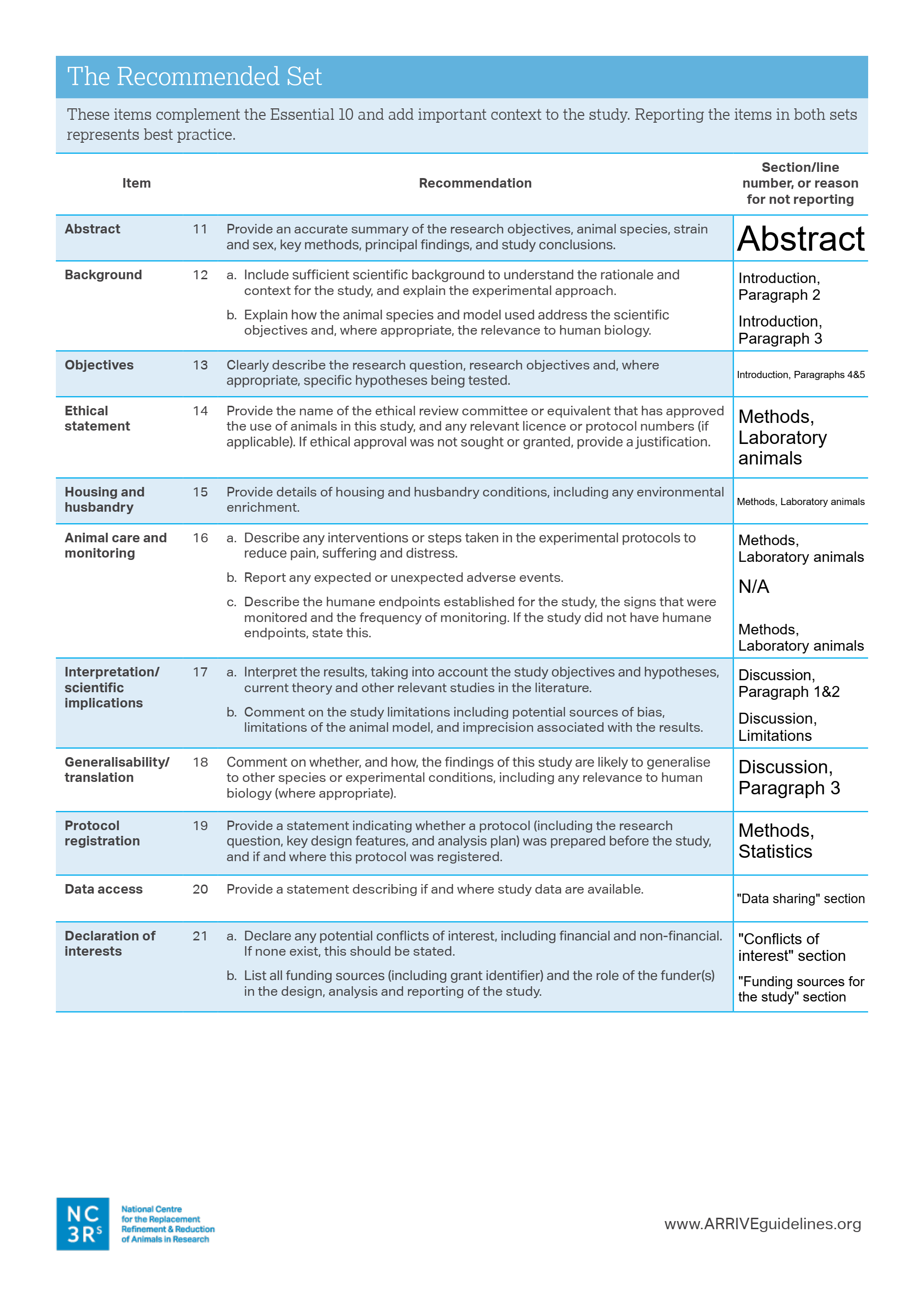
